# *Mycobacterium tuberculosis* cording in the cytosol of live lymphatic endothelial cells

**DOI:** 10.1101/595173

**Authors:** Thomas R. Lerner, Christophe J. Queval, Rachel P. Lai, Matthew Russell, Antony Fearns, Daniel J. Greenwood, Lucy Collinson, Robert J. Wilkinson, Maximiliano G. Gutierrez

**Author notes:** These authors contributed equally to the work (Thomas R. Lerner, Christophe J. Queval).

## Abstract

The ability of *Mycobacterium tuberculosis* to form serpentine cords is intrinsically related to its virulence, but specifically how *M. tuberculosis* cording contributes to pathogenesis remains obscure. We show that several *M. tuberculosis* clinical isolates form intracellular cords in primary human lymphatic endothelial cells (hLEC) *in vitro* and also in the lymph nodes of patients with tuberculosis. We identified via RNA-seq a transcriptional programme in hLEC that activates cellular pro-survival and cytosolic surveillance of intracellular pathogens pathways. Consistent with this, cytosolic access of hLEC is required for intracellular *M. tuberculosis* cording; and cord formation is dependent on the *M. tuberculosis* ESX-1 type VII secretion system and the mycobacterial lipid PDIM. Finally, we show that *M. tuberculosis* cording is a novel size-dependent mechanism used by the pathogen to evade xenophagy in the cytosol of endothelial cells. These results provide a mechanism that explains the long-standing association between *M. tuberculosis* cording and virulence.

## Introduction

*Mycobacterium tuberculosis* is one of the most successful bacterial pathogens of humankind and still constitutes a global health challenge (WHO, 2017). A striking phenotype of *M. tuberculosis* growing in nutrient broth is the ability of this pathogen to form serpentine cords, a morphological observation originally described by Robert Koch (Koch, 1882). This cording phenotype is intimately associated with virulence and immune evasion (Glickman et al., 2000). The first morphological descriptions of *M. tuberculosis* growth in liquid and solid media described a distinct ability of tubercle bacilli to form large and elongated structures by Middlebrook, Dubos and Pierce in the mid-1940s (Middlebrook et al., 1947). Cording is a complex phenotype involving many mycobacterial factors including lipids such as the “cord-factor” glycolipid trehalose dimycolate (TDM) (Hunter et al., 2006a; Hunter et al., 2006b; Indrigo et al., 2002) and a series of chemical modifications such as cyclopropanation of mycolic acids in the cell wall (Glickman et al., 2000).

Similar cording has been reported in other pathogenic mycobacteria, primarily in liquid media or extracellularly in various cell and organism models of infection. In zebrafish, *M. abscessus* released from apoptotic macrophages grows extracellularly, forming cords (Bernut et al., 2014). It is postulated that apoptosis of infected macrophages is a key event in the release of extracellular bacteria and subsequent initiation of cord formation. There are, however, a few reports showing that cording can also occur intracellularly. In 1928, Maximow and co-workers first reported intracellular cording in tissue culture (Maximow, 1928). In 1957, Shepherd studied this phenomenon in HeLa cells and found that only fully virulent *M. tuberculosis* strains formed cords. Moreover, Ferrer and co-workers (Ferrer et al., 2009) showed that an attenuated mutant of *M. tuberculosis* formed cords in fibroblasts.

Overall, extracellular cording has been shown in mycobacteria to be anti-phagocytic and to be a trigger of extracellular trap formation in macrophages (Bernut et al., 2014; Kalsum et al., 2017; Wong and Jacobs, 2013). Although proposed as a virulence mechanism, this does not explain why an intracellular pathogen such as *M. tuberculosis* would prefer to replicate in cords in the relatively nutrient poor extracellular space to avoid phagocytosis.

Bacterial xenophagy is the process that regulates the removal of cytosolic bacteria after damage to phagosomal membranes during selective macroautophagy (Galluzzi et al., 2017). This pathway constitutes one of the first cell autonomous defence pathways against intracellular pathogens (Deretic and Levine, 2009; Gutierrez et al., 2004). A fraction of the *M. tuberculosis* population damage phagosomes to access the cytosol and are subsequently recognised by autophagic adaptors and the xenophagy machinery. This process targets *M. tuberculosis* into autophagosomes and thus the lysosomal degradation pathway (Watson et al., 2012). Whereas there is a large body of literature demonstrating autophagy as an anti-mycobacterial pathway (Deretic et al., 2009), recent evidence shows that *M. tuberculosis* can eventually block the fusion of autophagosomes with lysosomes (Lerner et al., 2016; Romagnoli et al., 2012) and in mice, *M. tuberculosis* can evade autophagic responses *in vivo* (Kimmey et al., 2015).

*M. tuberculosis* mostly infects macrophages although there is compelling evidence that a minor proportion of *M. tuberculosis* is found infecting various non-myeloid cells in the lungs and lymph nodes *in vivo* (Ganbat et al., 2016; Lerner et al., 2015; Nair et al., 2016; Randall et al., 2015). The role that these *M. tuberculosis* subpopulations play in TB pathogenesis in different cell types (e.g. immune vs non-immune) is unclear. We previously showed in extrapulmonary tuberculosis that a subpopulation of *M. tuberculosis* is found in human lymphatic endothelial cells (hLEC) in lymph node biopsies and these cells could represent a reservoir for *M. tuberculosis* in infected patients (Lerner et al., 2016).

Here we discovered that *M. tuberculosis* forms large intracellular cords consisting of up to thousands of individual bacteria arranged end-to-end in hLEC *in vitro* and in biopsies of tuberculosis patients. Intracellular cording is common to all tested clinical isolates and ‘virulent’ lab strains of wild-type *M. tuberculosis* that had not lost the ability to produce phthiocerol dimycocerosates (PDIMs) during laboratory sub-culturing. We identified a transcriptional signature from the host consistent with *M. tuberculosis* membrane damage and escape from the phagosome into the cytosol and used correlative light electron microscopy (CLEM) to determine that intracellular cords are formed of chains of individual *M. tuberculosis* which are only present in the host cell cytosol. *M. tuberculosis* mutants lacking ESX-1 or PDIMs that cannot access the cytosol are incapable of cording unless co-cultured with wild-type bacteria to ‘smuggle’ them from a shared phagosome into the cytosol. Finally, we show that cords are devoid of endosomal, phagosomal and autophagosomal cellular markers and are formed from bacteria that successfully evaded p62-dependent xenophagy. Our results argue that cording represents an intracellular immune evasion strategy and then, once extracellular, anti-phagocytic. Our data also show that when growing, there is a size-dependent effect where bacteria are too large to be recognised by xenophagy. These results provide evidence for a bacterial pathogen size-dependent mechanism of xenophagy avoidance.

## Results

### *M. tuberculosis* forms extensive intracellular cords in hLECs

By monitoring GFP-expressing *M. tuberculosis* H37Rv (GFP-*M. tuberculosis*) replication in hLECs *in vitro* at different time points after infection, we observed a striking ability of *M. tuberculosis* to form distinctive intracellular cords (**Fig. 1a**). 3D confocal imaging confirmed these cords to be intracellular rather than on the cell surface (**Fig. 1b**). To quantitatively and accurately measure intracellular *M. tuberculosis* cording, we used the maximum feret diameter which describes the distance between the two furthest extremities of the cord (explained in **Supplementary Fig. 1**). Intracellular *M. tuberculosis* cords were very long, particularly after 72 h of infection, with a length of up to ∼150 μm (**Fig. 1c**). Intracellular cording was also observed in a human type II alveolar epithelial cell line (A549) although less prominent than in hLEC, likely due to the A549 cells themselves being smaller than hLEC (**Fig. 1d**). Importantly, intracellular cord formation was present not only in the lab-adapted strain H37Rv but also when hLEC were infected with any of three clinical isolates representing *M. tuberculosis* lineages (**Fig. 1e**). The cords were also present in lymph nodes of extrapulmonary TB patients (**Fig. 1f**). We observed that in Ziehl-Neelsen stained lymph nodes with TB granulomas, intracellular bacterial cords were present in cells with pleiotropic morphologies, including endothelial-like morphology (**Fig. 1f**). To confirm these observations, sections were stained for the lymphatic endothelial marker podoplanin (PDPN) and *M. tuberculosis* (Lerner et al., 2016). Despite only few LEC are infected with *M. tuberculosis*, the intracellular cording phenotype was associated with LEC in lymph node biopsies and the size of these cords ranged from 4 to 21 μm (**Fig. 1f**). Thus, *M. tuberculosis* has the ability to cord intracellularly in primary hLEC *in vitro* and in human lymph nodes of TB patients.

**Fig. 1:**
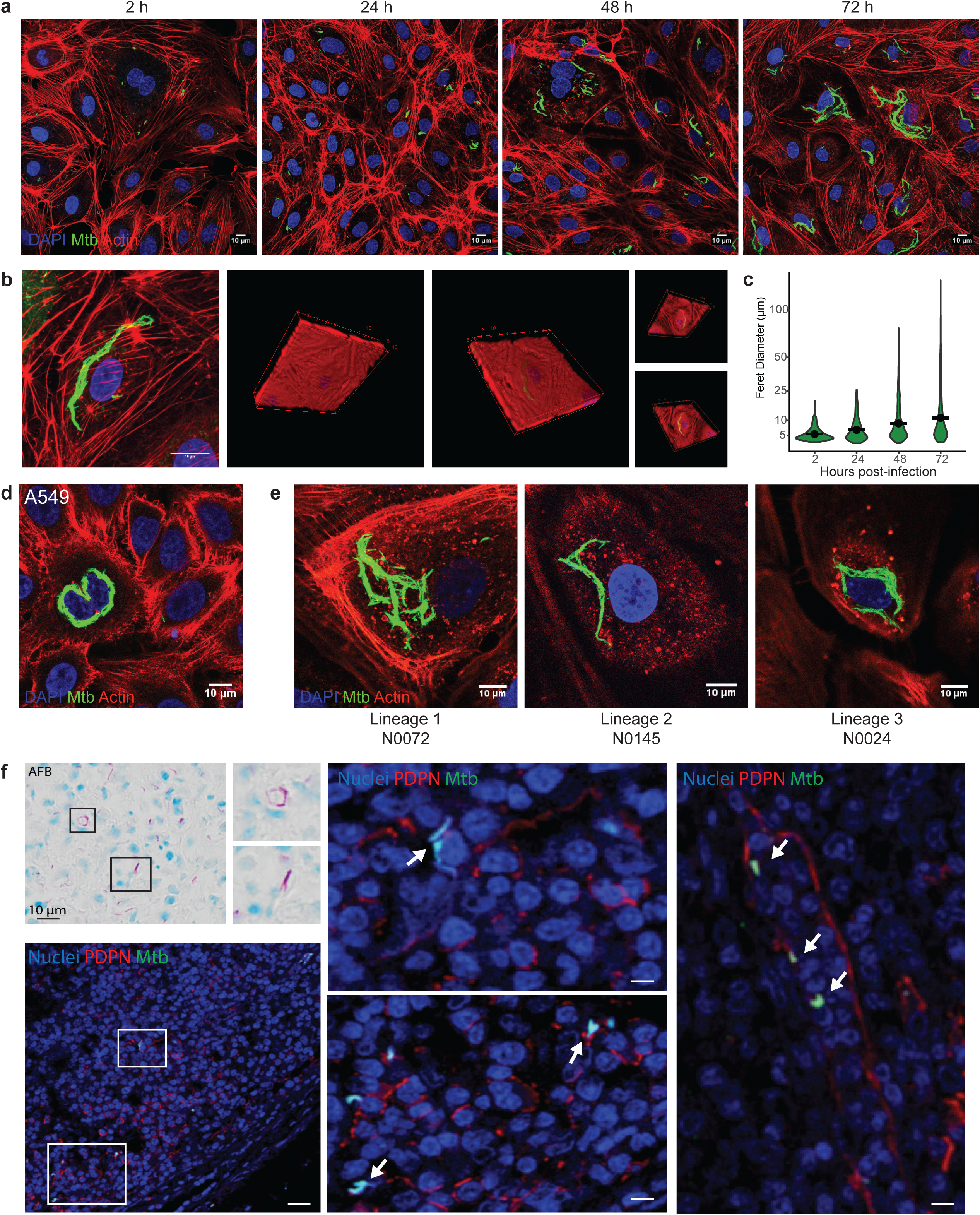
*M. tuberculosis* forms large intracellular cords *in vitro* and *in vivo*. **(a)** Images of primary human lymphatic endothelial cells (hLEC) infected with GFP expressing *M. tuberculosis* for 2-72 h. Over time, *M. tuberculosis* grows and forms large intracellular cords. Nuclei are stained with DAPI (blue) and F-actin is stained by rhodamine phalloidin (red). Scale bars are 10 μm. **(b)** 3D reconstruction of Z-stacks taken of an intracellular cord from (A). Various angles are shown to confirm that the cord is completely encapsulated within the host cell. Scale bar is 10 μm. **(c)** Measurement of the intracellular cords over time in hLEC using the Feret diameter (see **Supplementary Fig. 1**) showing that the cords elongate over time up to a maximum of 145 μm. The number of bacterial clusters analysed are: 418 (2h), 233 (24h), 814 (48h), 618 (72h) **(d)** Image of A549 cells infected with *M. tuberculosis*-EGFP for 72 h showing an intracellular cord looping around the nucleus. Nuclei are stained with DAPI (blue) and F-actin is stained with rhodamine phalloidin (red). Scale bar is 10 µm. **(e)** Intracellular cord formation after 72 h was also observed in hLEC infected with representative strains from three other *M. tuberculosis* lineages: N0072 (lineage 1), N0145 (lineage 2), N0024 (lineage 3). **(f)** Tissue section of a granuloma present in a human lymph stained for acid fast bacilli (AFB). Zoomed region shows association of *M. tuberculosis* cords with cells (black arrows). Representative histological sections from human patients after lymph node tissue resection surgery were stained for podoplanin (PDPN), *M. tuberculosis* and nuclei (DAPI). Arrows indicate the presence of *M. tuberculosis* cords within PDPN+ cells. Scale bar is 1 mm.

### *M. tuberculosis* infection induces cytosolic surveillance of bacterial pathogens and pro-survival response in hLECs

To better understand the host cell response to the extensive *M. tuberculosis* cording in the cytosol, we performed RNA-seq analysis in uninfected and *M. tuberculosis*-infected hLEC at 48 h after infection when cords started to be prominent. Among the top ten statistically significant process networks induced by *M. tuberculosis* infection were inflammation and interferon signalling, antigen presentation, phagosome antigen presentation, and innate immune response (**Fig. 2a**). Notably, in addition to a strong pro-inflammatory response (**Fig. 2b**), we identified four additional pathways that were significantly up-regulated after infection (**Fig. 2b**). There was an upregulation of pathways that recognised cytosolic RNA and dsDNA with an upregulation of type I interferon. The pathways of cytosolic carbohydrate recognition as well as STING signalling were also upregulated suggesting a high level of membrane damage induced by *M. tuberculosis* (**Fig. 2b**). Importantly, RNA-seq identified a transcriptional signature consistent with anti-cell death or pro-survival pathways and antigen presentation (**Fig. 2c**). This pro-survival signature was unexpected based on our data on human primary macrophages (Lerner et al., 2017), although consistent with previous live-cell observations that active *M. tuberculosis* replication in primary hLECs was not associated with significant host cell death (Lerner et al., 2016). We confirmed by RT-qPCR that the expression of the pro-inflammatory cytokine IL-6 as well as the type I IFN responsive cytokines CXCL10 (IP10) and IFN-β were significantly upregulated after infection in hLECs (**Fig. 2d**). The pro-survival factors BCL2A1, EIF2AK2 and TNFAIP3 (A20) were also significantly up-regulated (**Fig. 2d**). Moreover, the cytosolic glycan sensing genes Galectin-3, Galectin-8, cGAS and the foreign DNA sensor ZBP1 were upregulated after infection (**Fig. 2d**). In the case of Gal-3, a high level of expression was observed already in uninfected cells (**Fig. 2d**). Thus, infection of hLECs with *M. tuberculosis* induced host pro-survival pathways and negative regulators of cell death to protect the niche in which bacteria proliferate. On the other hand, endothelial cells upregulated cytosolic surveillance of RNA, DNA and carbohydrates pathways to recognise *M. tuberculosis* in the cytosol.

**Fig. 2:**
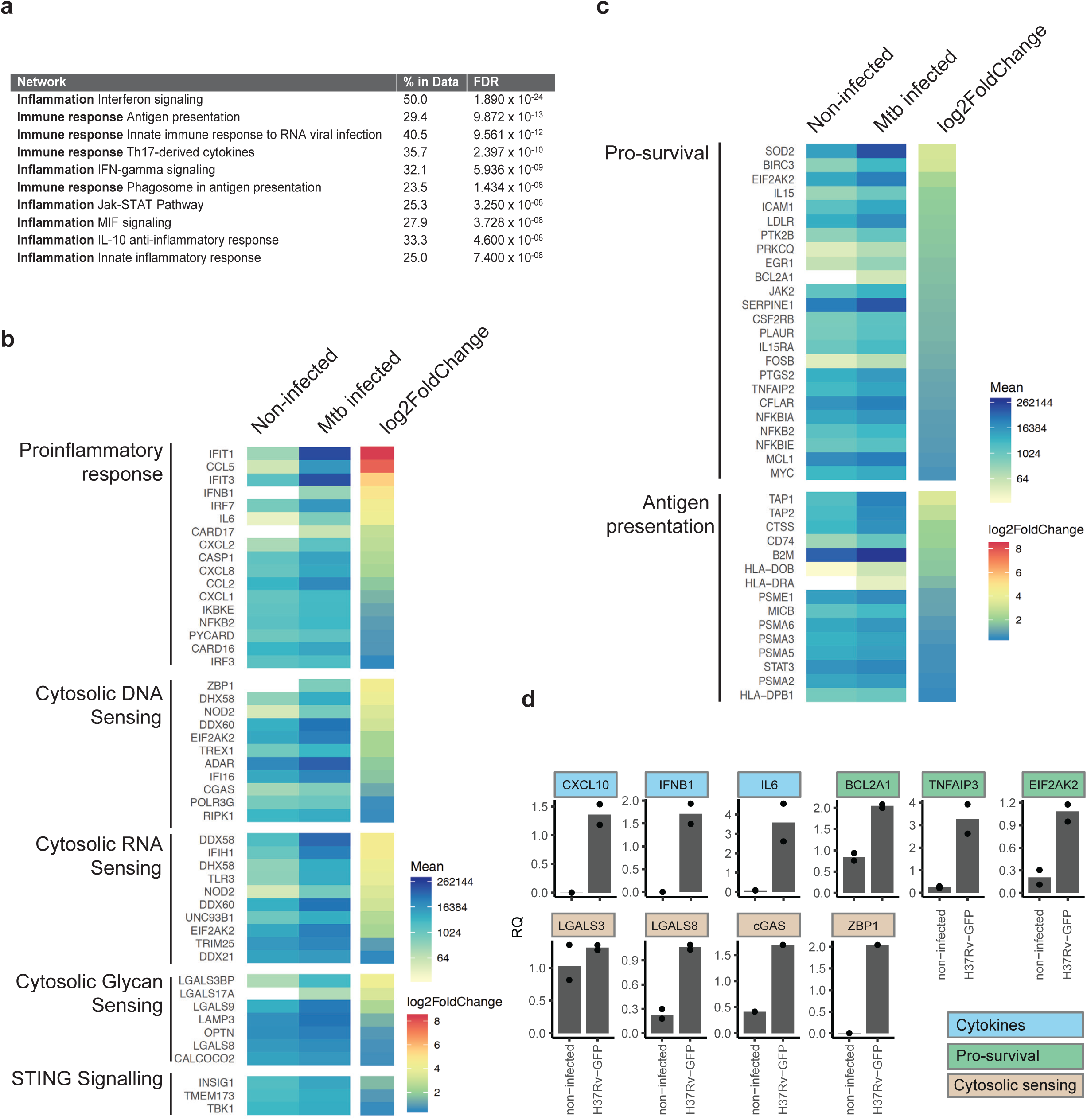
*M. tuberculosis* induces cytosolic surveillance and host pro-survival pathways. **(a)** Top 10 functional process analysis hits by false-discovery rate (FDR) of genes significantly upregulated in hLECs 48 h post-infection, indicated by RNAseq. ‘% in Data’ indicates the % of genes in the annotation group that were significantly upregulated in the analysis. **(b)** Heatmap of significantly upregulated (padj < 0.05) genes 48 h post infection grouped by sensing pathway reveal an induction of pro-inflammatory, DNA, RNA and glycan sensing pathways and **(c)** genes involved in antigen presentation and the negative regulation of cell death. (**d**) qPCR confirmation of key infection-response pathways 48 h post infection.

### *M. tuberculosis* intracellular cording requires both RD1 and PDIM

We next sought to understand *M. tuberculosis* factors that contributed to the intracellular cording phenotype in hLEC. We have previously shown that the ESX-1 secretion system, encoded in the RD1 genomic region, and the cell wall lipid phthiocerol dimycocerosate (PDIMs) are required for intracellular replication of *M. tuberculosis* in hLEC (Lerner et al., 2016; Lerner et al., 2018). Infection with the *M. tuberculosis* ΔRD1 mutant that lacks the ESX-1 secretion system was not able to form cords but instead exhibited smaller clumps of bacteria sometimes with a mesh-like appearance (**Fig. 3a**). The phenotype of the *M. tuberculosis* mutant lacking PDIM also presented a clumpy mesh-like phenotype with an increased number of individual bacteria that were not organised in cords (**Fig. 3a**). *M. tuberculosis* mutants lacking either the ESX-1 secretion system or the virulence-related lipid PDIM (Astarie-Dequeker et al., 2009) failed to cord intracellularly (**Fig. 3b**). The lack of cording observed with the RD1 mutant was not due to the reduced bacterial burden, since increasing the multiplicity of infection did not increase cord formation although significant bacterial growth was observed (**Fig. 3c, d, e, f**). Moreover, we found that the up-regulation of some genes in hLEC after infection (**Fig. 2b**) such as interferon-beta (IFN-β) or interleukin-6 (IL-6) was RD1 and PDIM dependent (**Supplementary Fig. 2**). For other genes, ESX-1 and PDIM seem to play a suppressive role, suggesting that other Mtb factors are involved in the activation of immune pathways. Altogether, in hLEC, the ability of *M. tuberculosis* to form intracellular cords requires both the ESX-1 system and the lipid PDIMs.

**Fig. 3:**
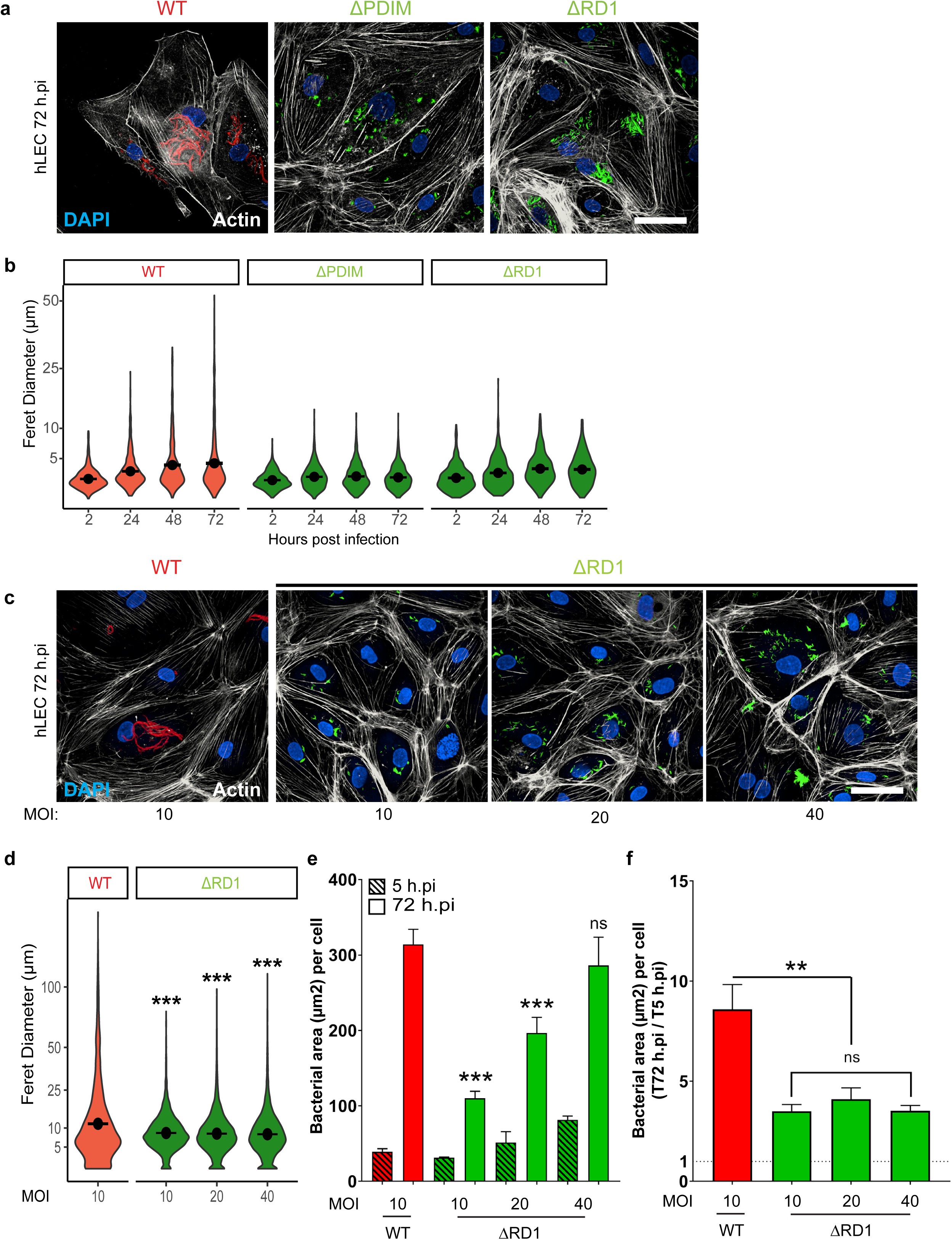
Cord formation requires both ESX-1 and PDIM. **(a)** hLEC were infected with RFP-expressing *M. tuberculosis* WT, GFP-expressing *M. tuberculosis* ΔPDIM or E2-Crimson-expressing ΔRD1 for 72 h at a MOI of 10, fixed and stained for F-actin with AF633 or AF488-phalloidin. Either deleting PDIM or the RD1 locus abolished cord formation. WT bacteria (red), ΔPDIM and ΔRD1-bacteria (green), F-actin (white) and nuclei (blue). Scale bar is 50 µm. **(b)** Feret diameter measurements from three independent experiments were plotted. For each condition tested, the number of bacterial clusters analysed is between 600 and 1,200 **(c)** hLEC were infected for 72 h with RFP-expressing *M. tuberculosis* WT at a MOI of 10, or with E2-Crimson-expressing *M. tuberculosis* ΔRD1 at a MOI of 10, 20 or 40. WT bacteria (red), ΔRD1-bacteria (green), F-actin (white) and nuclei (blue). Scale bar is 50 µm. **(d)** Feret diameter measurements from images in (C) from two independent experiments were plotted. The number of bacterial clusters analysed are: 3,960 for WT and 6,470, 9,472, 11,759 for ΔRD1 at MOI:10, 20 and 40, respectively. (**e**) Quantification of the bacterial load per cell, expressed in bacterial area (µm^2^) per cell, following the uptake (5h.pi) and 72h post infection. (**f**) Intracellular bacterial growth after 72h infection, expressed by the ratio bacterial area per cell 72h.pi/5h.pi. Values > 1 represent the bacterial growth.

### *M. tuberculosis* intracellular cords are localised in the cytosol

Given that at least two critical *M. tuberculosis* virulence-associated factors that contribute to cytosolic localisation were required for intracellular cording and the significant upregulation of cytosolic pathogen surveillance during cording, we next sought to define the subcellular compartment within which *M. tuberculosis* cords were localised in hLECs. By using a correlative imaging approach (correlative light and electron microscopy, CLEM), we determined that *M. tuberculosis* intracellular cords were localised in the cytosol of hLEC in long structures that (in this example) looped around the host cell nucleus (**Fig. 4a**). In contrast, small groups of *M. tuberculosis* containing relatively low numbers of individual bacteria were localised in a membrane-bound compartment (**Fig. 4b**) as reported before (Lerner et al., 2016). The cords are usually formed of a bundle of several parallel chains of *M. tuberculosis* (**Fig. 4c**) and therefore a single cord can consist of (up to) thousands of individual bacteria. Interestingly, the volume of 25 individual bacteria from a cord compared to 25 from a membrane bound compartment (non-cord, displayed as coloured reconstructions) as measured by three-dimensional serial block face (3D SBF) CLEM was significantly lower (**Fig. 4d**). This confirmed that the cords did not consist of *M. tuberculosis* which had become abnormally long/filamentous. We concluded that *M. tuberculosis* intracellular cording occurs in the cytosol of hLEC and that cytosolic *M. tuberculosis* cords are composed of hundreds or thousands of individual mycobacteria that are smaller than bacteria contained in membrane-bound compartments.

**Fig. 4:**
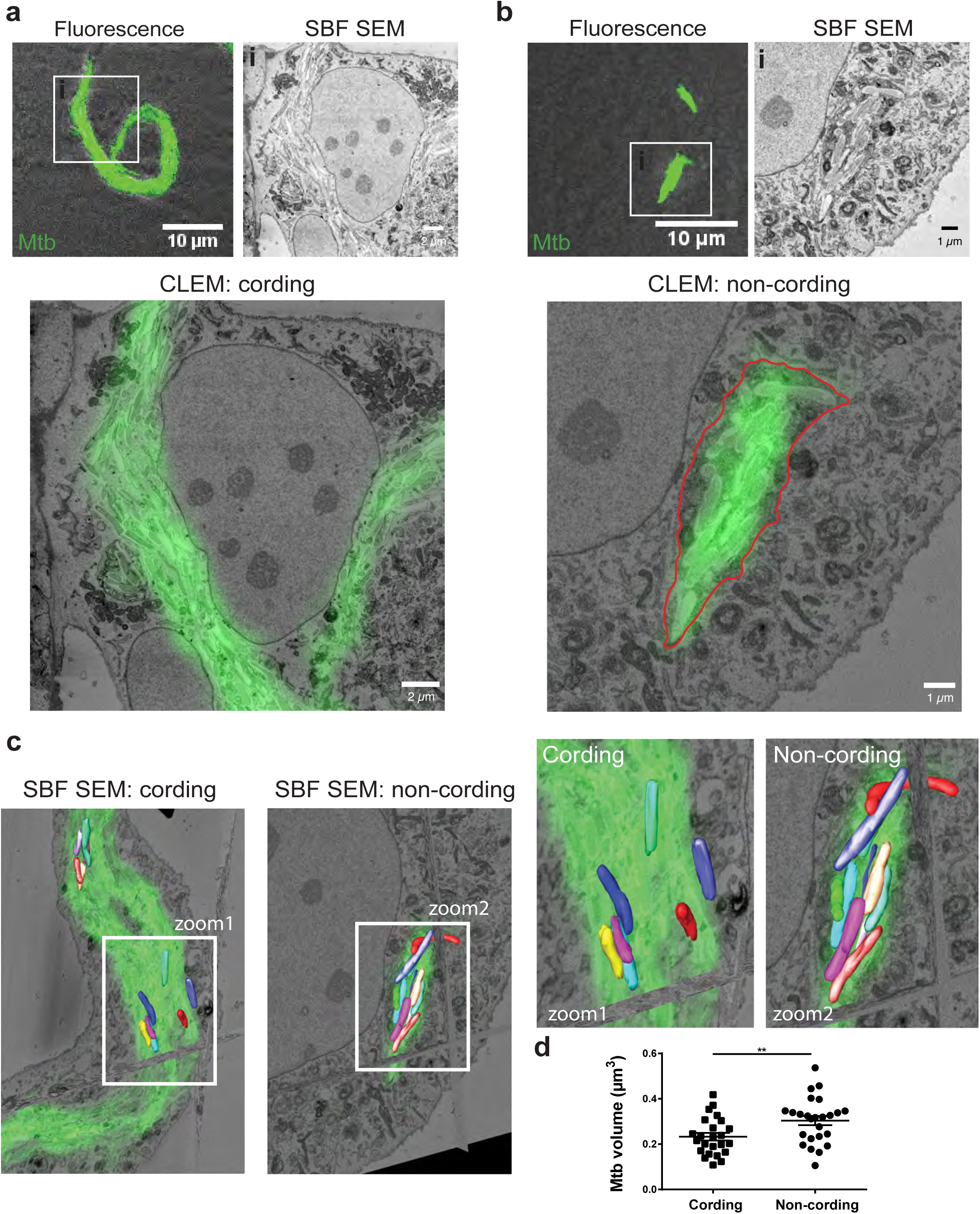
Intracellular cords are localised in the host cell cytosol and consist of chains of *M. tuberculosis* of a small size. **(a-b)** Correlative light electron microscopy (CLEM) images of hLEC infected with *M. tuberculosis*-GFP. Top left subpanel shows the light microscopy images, with the corresponding electron microscopy images in the top right subpanel. The larger subpanels show a composite of the fluorescence overlaid onto the electron microscopy. **(a)** *M. tuberculosis* intracellular cord, without any encapsulating host membrane, indicating that it is present in the cytosol. **(b)** *M. tuberculosis* encapsulated in a membranous compartment, as a control for confirming membrane preservation due to the sample preparation. Host cell membrane is highlighted in red. Scale bars are 10 μm. **(c-d)** To quantify the volume of *M. tuberculosis*, individual bacteria were manually segmented from slices of SBF SEM images and 3D reconstructions of selected bacteria were made (coloured rods), using 3dmod. **(c)** Representative reconstructions are shown, with corresponding fluorescence highlighted (matched manually with the corresponding SBF SEM slice in Z, and then aligned in xy with TurboReg in Fiji). Dataset dimensions; left panel 8.7 × 8.7 × 50 nm pixels, 71.3 × 71.36 × 1 µm in xyz; right panel 6.3 × 6.3 × 50 nm pixels; 51.6 × 51.6 × 2.75 µm in xyz. **(d)** The volume of each bacterium reconstruction from two independent sample datasets was calculated in 3dmod, and a comparison between those in a membrane bound compartment and those in an intracellular cord was made. The data show that the cords are formed from *M. tuberculosis* which are significantly smaller. Student’s t-test; ** = p <0.01.

### Access to the cytosol is required for *M. tuberculosis* replication and intracellular cording

The localisation of *M. tuberculosis* cords suggested that the cytosol represents a permissive environment for *M. tuberculosis* replication, thus we tested if the RD1 mutant of *M. tuberculosis* that is mostly localised in membrane-bound compartments could replicate and form intracellular cords if forced to access the cytosol. To achieve that, we performed a series of co-infection experiments combined with CLEM. As shown before, in RFP-*M. tuberculosis* H37Rv WT single infection of hLEC, RFP-*M. tuberculosis* WT formed prominent intracellular cords whereas single infection with E2-Crimson-*M. tuberculosis* ΔRD1 or GFP-*M. tuberculosis* ΔPDIM did not show cording (**Fig. 5a**). Strikingly, if hLEC are co-infected with RFP-*M. tuberculosis* WT and with E2-Crimson-*M. tuberculosis* ΔRD1 or GFP-*M. tuberculosis* ΔPDIM, the *M. tuberculosis* mutants lacking either ESX1 or PDIM were now able to clearly form intracellular cords (**Fig. 5a**). Consistent with these observations, the feret diameter of E2-Crimson-*M. tuberculosis* ΔRD1 or GFP-*M. tuberculosis* ΔPDIM in co-infected cells was similar to RFP-*M. tuberculosis* WT whereas GFP-*M. tuberculosis* ΔRD1 or ΔPDIM alone had low feret diameter measurements (**Fig. 5b**). Importantly, in co-infected cells, both the *M. tuberculosis* ΔRD1 or ΔPDIM were able to replicate more efficiently (**Fig. 5c**). By CLEM, we confirmed that the RFP-*M. tuberculosis* WT was localised in the cytosol and defined at the ultrastructural level that the cords formed by GFP-*M. tuberculosis* ΔRD1 in co-infected cells were now localised in the cytosol (**Fig. 5d, e**). Altogether, *M. tuberculosis* replicates in the cytosol of hLEC forming long intracellular cords; moreover, because bacteria that normally do not access the cytosol such as the *M. tuberculosis* ΔRD1 mutant were able to do so when forced into the cytosol, this indicates that the effect of ESX-1 on intracellular cording is mediated by access to the cytosol.

**Fig. 5:**
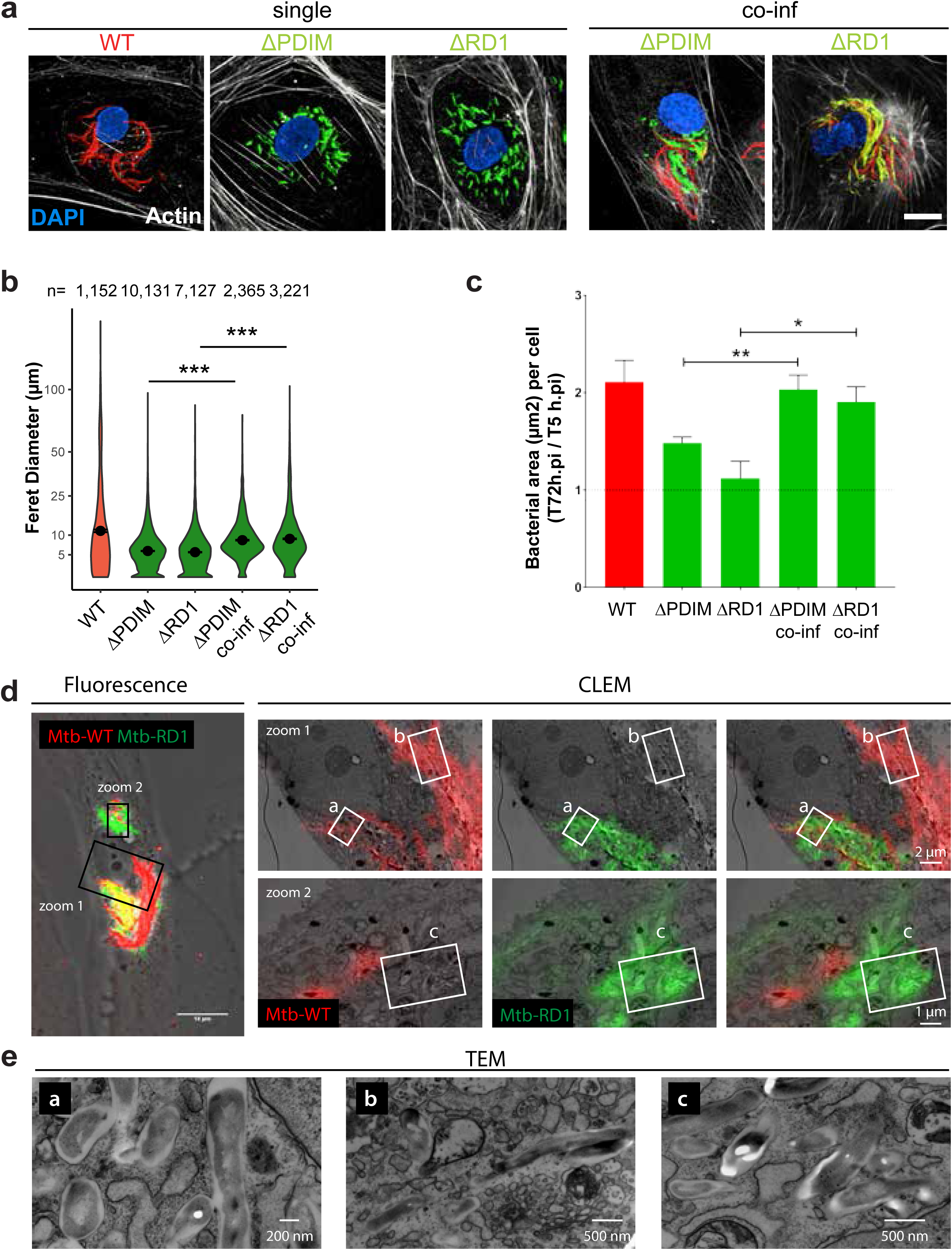
Access to the cytosol is required for *M. tuberculosis* intracellular cording. **(a)** hLEC were infected for 72 h with *M. tuberculosis* WT-RFP (red), *M. tuberculosis*-ΔPDIM-GFP (green), *M. tuberculosis*-ΔRD1-E2-Crimson (green) either individually or as a co-infection WT-RFP/ΔPDIM-GFP or WT-RFP/ΔRD1-E2-Crimson. Cells were then fixed and stained for F-actin with AF633 or AF488-phalloidin (Both visualized in white) and DAPI (blue). Scale bar is 10 µm. The images show that during single infection, *M. tuberculosis* WT exhibits intracellular cording, whereas *M. tuberculosis* ΔPDIM or ΔRD1 do not. However, in the co-infected sample, both *M. tuberculosis* ΔPDIM and ΔRD1 were able to form intracellular cords. **(b)** Feret diameter measurements from images in **(a)** were plotted. n represent the number of bacterial clusters analysed. **(c)** Intracellular bacterial growth after 72h infection, expressed by the ratio bacterial area per cell 72h.pi/5h.pi. Values > 1 represent the bacterial growth. **(d-e)** Co-infected hLEC samples were processed for correlative light electron microscopy (CLEM) to confirm at the ultrastructural level that *M. tuberculosis* ΔRD1-GFP cords were indeed present in the cytosol (e; magnifications of regions indicated in d).

### Intracellular cords form from bacteria that evaded xenophagy

Because *M. tuberculosis* intracellular cords formed in the cytosol and induced a pathogen cytosolic recognition signature in hLEC associated with xenophagy (Watson et al., 2012a), we investigated whether *M. tuberculosis* cords were targeted by selective autophagy. In hLEC, *M. tuberculosis* targeting via selective autophagy is PDIM dependent (Lerner et al., 2018) and entirely RD1 dependent (**Supplementary Fig. 2**) suggesting that in hLECs, xenophagy primarily recognises mycobacteria that access the cytosol, the intracellular location for *M. tuberculosis* cording. Thus, we investigated whether cording vs non-cording populations of *M. tuberculosis* were recognised by the selective autophagy machinery. Notably, when we co-labelled ubiquitin and p62 in cord-containing cells, we found that both markers selectively associated with only small bacterial groups and not *M. tuberculosis* cords (**Fig. 6a**). Strikingly, large *M. tuberculosis* cords (as defined by having a feret diameter of greater than 10 µm), were devoid of the selective autophagy markers ubiquitin, p62, Galectin-8, NDP52 and LC3B as well as the late endosomal/lysosomal markers LAMP-2 and cathepsin D (**Fig. 6b**). In contrast, we found that, whereas none of the markers analysed localised to the cords, some of the markers localised to a single or small group/clump of intracellular *M. tuberculosis* with a lower feret diameter (**Fig. 6b**). These data indicated that, although large and long *M. tuberculosis* cords were present in the cytosol, these were not recognised by xenophagy. Consistent with the cords being negative for the autophagy-related host-cell markers tested, live cell imaging in hLECs expressing RFP-p62 revealed that the intracellular cords form from bacteria which have either completely evaded p62-positive compartments as a readout of autophagic targeting (**Fig. 6c, Movie S1**) or which have initially been growth-restricted in a p62-positive state (**Fig. 6d**) but subsequently became p62-negative, where this process can also cycle several times (**Fig. 6e, Movie S2**). Crucially, the *M. tuberculosis* cords only ever form once the bacteria lost p62 (**Fig. 6e, Movie S3**) suggesting that cording is a consequence of avoiding an autophagic state or that cord formation blocks autophagic targeting, potentially by being too large to encapsulate and recapture from the cytosol.

**Fig. 6:**
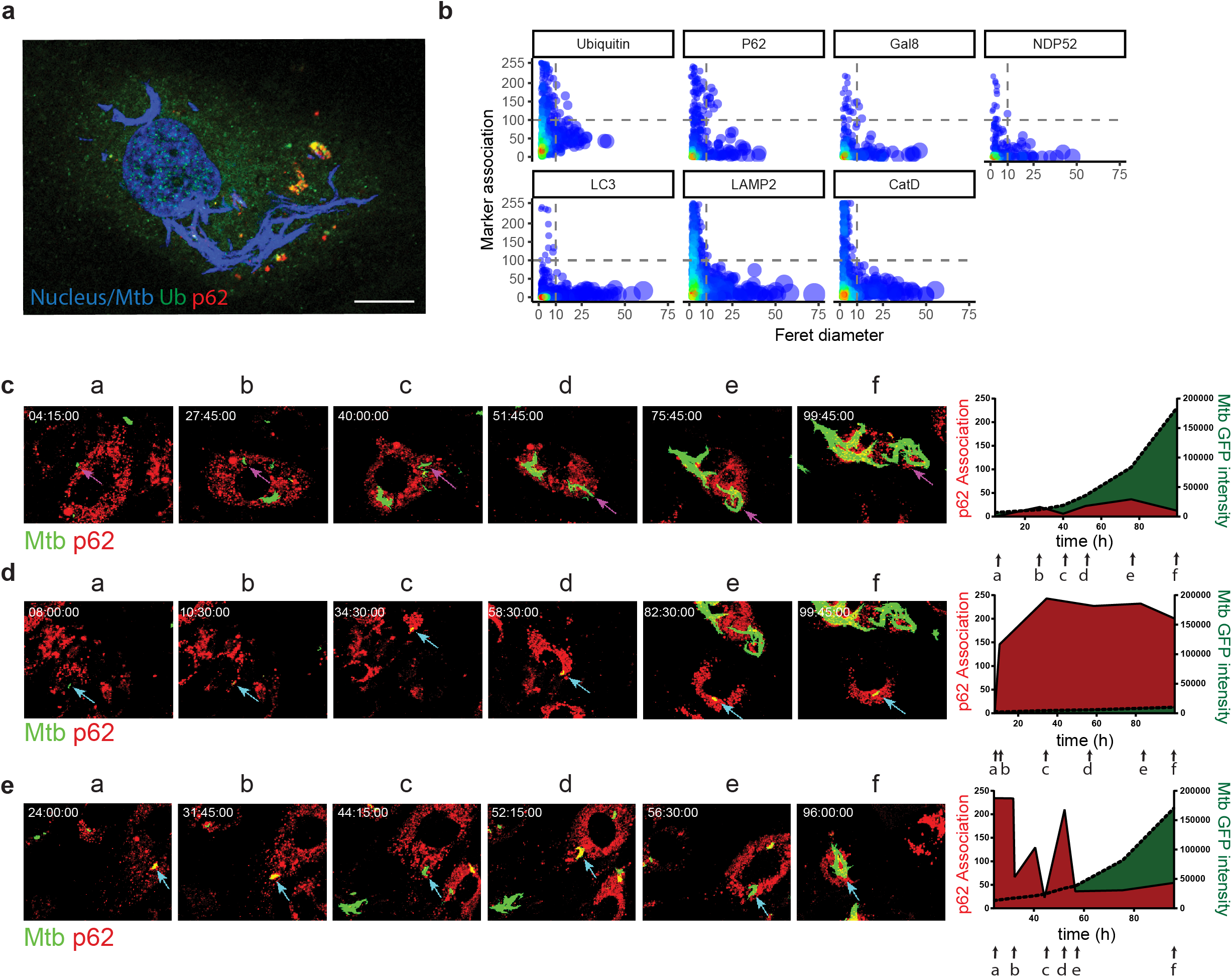
Intracellular cords form from individual bacteria that evade or escape selective autophagy. **(a)** Representative image of hLEC infected *M. tuberculosis* WT-EGFP (blue) for 72 h and stained for the autophagy adaptor p62 (red) and the autophagy receptor ubiquitin (Ub) (green). Cell nuclei are stained with DAPI (blue). Scale bar is 10 μm. **(b)** Intracellular markers of autophagy, pathogen sensing were assessed for their association to intracellular cords 72 h post infection. Particles with a feret diameter greater than 10 μM were considered cords, and a marker association score above 100 was considered positive. Points correspond to individual bacterial particles, with diameters scaled to the Feret diameter of the particles. Colour indicates density of points, with red indicating high density and blue low density. **(c-e)** Live cell imaging of hLEC expressing p62-RFP (red) infected with *M. tuberculosis*-GFP (green) for 115 h. Imaging started 15 min after addition of the bacteria to the cells. Snapshots from the movies (Movies S1-3) are shown, with the timepoint displayed above in hh:mm:ss format. Scale bars are 10 μm. **(c)** The pink arrow tracks an example of an intracellular cord forming from a single bacterium, which never interacts/associates with p62. **(d)** The blue arrow tracks an example of an individual *M. tuberculosis* bacterium becoming associated to p62 throughout which leads to restriction of growth. **(e)** The blue arrow tracks an example of *M. tuberculosis* associating and dissociating with p62 multiple times. Only after p62 association ceased completely, cord formation started. **(c-e, right hand panel)** ImageJ quantification of the GFP intensity and the p62-RFP association of the arrowed bacteria over time. Letters a-f refer to the snapshots in (**c-e**).

## Discussion

Since the identification of *M. tuberculosis* as the etiologic agent of human TB, the phenomenon of cording has attracted significant interest because of its association with virulence and infection *in vivo*. Whereas there are many studies that implicate cording as a mechanism to subvert phagocytosis, there is little evidence that in the infected host, *M. tuberculosis* can freely replicate and cord in the extracellular milieu to avoid phagocytosis. We show here that *M. tuberculosis* intracellular cords are a size-dependent mechanism of evasion of endothelial host cell intracellular innate immune defences such as xenophagy. We postulate that cords are linked to virulence because bacteria can replicate to a large extent intracellularly within non-immune cells in a protected environment until nutrients are exhausted and space to grow is limited. Mycobacteria are then released into the extracellular milieu where large cords can block phagocytic uptake, allowing dissemination of *M. tuberculosis*. This is similar to the extracellular cords that form in the *M. absessus* infected zebrafish model where cords are too large to be phagocytosed and therefore facilitate immune evasion (Bernut et al., 2014). We determined that intracellular cording is a result of evading the host cell defences and allows vast numbers of bacteria to proliferate, only being stopped by physical space and eventually leading to the cell being compromised and cords disseminating, which are too large for phagocytosis by macrophages and/or neutrophils.

High burdens of cytosolic bacteria without induction of host cell death was surprising and suggested that human endothelial cells respond differently to infection that in human primary macrophages (Lerner et al., 2017). Several pathological studies have shown that while some bacilli produce massive tissue damage, especially in the lung, others persist in many tissues with no gross evidence of damage (Hunter et al., 2016). We propose that infection in macrophages tends to induce necrotic cell death whereas endothelial cells are more resistant to cell death and permissive for *M. tuberculosis* growth. Consistent with the prolonged survival of *M. tuberculosis*-infected hLEC, there is an *M. tuberculosis*-induced transcriptional signature of cell death present but this is alongside the upregulation of several anti-cell death and pro-survival pathways. Our studies are consistent with early observations in HeLa cells that found that only fully virulent *M. tuberculosis* strains could cord, often filling the whole cell without causing cytotoxicity (Shepard, 1957). Finally, our data provide an explanation for the observation that endothelial cells are infected in patients with tuberculosis but the typical clinical symptoms of endothelial damage are not observed as in other infectious diseases.

Our study also sheds some light on the preferred site of replication of *M. tuberculosis* in endothelial cells. Our experiments clearly show that if bacteria access the cytosol, they can cord and replicate. This suggests that the environment in membrane bound compartments is restrictive and the cytosol highly permissive for bacterial replication and cording. It remains to be defined if that is the case for macrophages. Interestingly, in one study that investigated the localisation of *M. tuberculosis* in resected lungs of tuberculosis patients, prominent cords were observable within macrophages at the luminal side of the granuloma cavity (Kaplan et al., 2003). If our studies in human cells and tissue are reflected in mice remains to be determined, howeve, the evasion of xenophagy by intracellular cording might provide an explanation for the reported evasion of this pathway in the mouse model of tuberculosis (Kimmey et al., 2015).

What determines that a subpopulation of intracellular *M. tuberculosis* starts cording? It is possible that differential expression of *M. tuberculosis* secreted or cell-surface proteins cause differential recognition of cytosolic *M. tuberculosis* by the autophagy apparatus. Our data show that the previously reported ubiquitin-mediated autophagy process by which *M. tuberculosis* extracellular DNA/RNA is recognised by the cGAS/STING pathway (Watson et al., 2012b) is also activated in hLEC. Whether it is the bacteria themselves that are ubiquitinated or their compartment is uncertain. If *M. tuberculosis* retains its waxy cell wall in the cytosol it is unlikely that ubiquitination will play a major role in xenophagic targeting. We reason that if the bacteria themselves are being recognised, then why is only a subpopulation targeted to autophagy? What is different about them? We hypothesise that it is the ESX-1 mediated damaged membranes surrounding bacteria that are recognised, and if *M. tuberculosis* is in close proximity to this it will be ‘captured’ with it. This process may be cyclical, with *M. tuberculosis* then damaging the autophagic compartment to escape again. However, if *M. tuberculosis* can get away from the damaged membranes after cytosolic translocation, it may be able to evade autophagic capture. This is likely to occur for the majority of the *M. tuberculosis*, hence why only a relatively small population are targeted to autophagy. It is unlikely that dead bacteria or those that do not damage the phagosomal membrane will be targeted to autophagy because it is ESX-1 and PDIM dependent; these populations are thus likely to mature into phagolysosomes. Although the cording phenotype seems to be unique for pathogenic mycobacteria, it remains to be determined if other cytosolic pathogens also evades autophagy in a size-dependent manner as shown here.

### Experimental procedures

#### Cells

Primary hLEC taken from inguinal lymph nodes (ScienCell Research Laboratories, #2500) were cultured according to the manufacturer’s instructions up to passage 5 as described fully in (Lerner et al 2016). For confocal microscopy of fixed cells, 20,000 cells in 300 μl complete endothelial cell medium (ECM) (ScienCell Research Laboratories, #1001) were seeded onto 10 mm diameter #1.5 glass coverslips (Glaswarenfabrik Karl Hecht, #1001/10_15). For imaging destined for CLEM, 10,000 cells per dish (MatTek, #P35G-1.5-14-CGRD) in 500 μl ECM were seeded to achieve a confluence of 30-50% (thus allowing visualisation of the grid reference etched into the dish). For live cell imaging, 25,000 cells per dish in 500 μl ECM were seeded to achieve a confluence of >80% (thus limiting the cells’ movement away from the field of view). For electron microscopy, 200,000 cells per T25 flask were seeded in 5 ml ECM. For imaging with the automated confocal microscope Opera Phenix, 5,000 cells per well were seeded in 96 well plate (Cell Carrier 96 ultra, PerkinElmer). Type II alveolar epithelial A549 cells (ATCC) were cultured according to the manufacturer’s instructions. For confocal microscopy, 50,000 cells in 500 µl DMEM (Gibco) supplemented with 10% (v/v) heat inactivated foetal calf serum (FCS) were seeded onto 10 mm diameter #1.5 glass coverslips.

### *Mycobacterium tuberculosis* strains

This study used the following EGFP tagged strains as described previously (Astarie-Dequeker et al., 2009; Lerner et al., 2016; Lerner et al., 2018): *Mycobacterium tuberculosis* H37Rv-EGFP (*M. tuberculosis* WT), *M. tuberculosis* H37Rv-EGFP ΔRD1 (*M. tuberculosis* ΔRD1), *M. tuberculosis* H37Rv-EGFP ΔRD1::RD1 (*M. tuberculosis* ΔRD1::RD1). In this study, we refer to the *M. tuberculosis*-GFP WT strain as *M. tuberculosis* WT, the *M. tuberculosis*-GFP PMM100 strain as *M. tuberculosis* ΔPDIM. Additionally, we have used M. tuberculosis H37Rv-RFP (tagged with plasmid pML2570) and H37Rv-ΔRD1-E2-Crimson (tagged with plasmid pTEC19, which was a gift from Lalita Ramakrishnan (Addgene plasmid # 30178) (Takaki et al., 2013). The clinical isolates *M. tuberculosis* N0072-EGFP (Lineage 1), *M. tuberculosis* N0145-EGFP (Lineage 2), *M. tuberculosis* N0024-EGFP (Lineage 3) were obtained from Sebastien Gagneux (Basel, Switzerland). Mycobacteria were cultured in Middlebrook’s 7H9 broth medium (Sigma-Aldrich, #M0178) supplemented with 10% (v/v) Middlebrook OADC (BD Biosciences, #212351) and 0.05% (v/v) Tween80 (Sigma-Aldrich, #P1754) in 50 ml Falcon tubes at 37°C with rotation. Alternatively, mycobacteria were plated on petri dishes containing Middlebrook’s 7H11 agar medium (Sigma-Aldrich, #M0428) supplemented with 10% OADC and incubated at 37°C for 2-3 weeks until colonies appeared.

### Infection of hLEC with *M. tuberculosis*

A detailed infection protocol can be found in (Lerner et al., 2016). Briefly, *M. tuberculosis* cultures were grown to mid-exponential phase, washed twice with PBS, once with ECM medium, and then shaken with glass beads to break up bacterial clumps. *M. tuberculosis* were then resuspended in ECM medium and centrifuged at a slow speed to pellet any remaining clumps, but leaving individual bacteria in suspension. The OD_600_ of the bacterial suspension was measured and then added to hLECs at a theoretical multiplicity of infection (MOI) of 10 in ECM medium. Infection was for five hours and was followed by two PBS washes to remove any uninfected *M. tuberculosis*. The infected cells were incubated usually for 2-72 h but up to 7 days for live cell imaging. For experiments requiring co-infection of two *M. tuberculosis* strains, we used strains tagged with different colours to distinguish between them (RFP, EGFP or E2-Crimson). These strains were prepared individually using the above method, and only mixed just prior to hLEC infection (at an MOI of 5 each, to achieve a total MOI of 10).

### Indirect immunofluorescence

An extended method can be found in (Lerner et al 2016). In summary, infected hLEC on coverslips were fixed with 3% methanol-free paraformaldehyde (Electron Microscopy Sciences, #15710) in PBS for 24 h. Coverslips were quenched with 50 mM NH_4_Cl (Sigma-Aldrich, #A9434) and then permeabilised with 0.01% saponin (Sigma-Aldrich, #84510) 1% BSA (Sigma-Aldrich, #A3912) in PBS. The cells were washed with PBS and then 30-50 μl of the primary antibody (diluted in PBS with 0.01% saponin, 1% BSA) was added onto the coverslips for one to two hours at room temperature (detailed in Table 1). Following this, three PBS washes preceded addition of the secondary antibody (diluted in the same way as the primary antibody) for one hour at room temperature. The coverslips were again washed three times in PBS, before an optional staining step for F-actin using a 1:250 dilution of either rhodamine phalloidin (Biotium, #00027), Alexa Fluor 633-phalloidin (Life Technologies, #A22284) or Alexa Fluor 488-phalloidin (Life Technologies, #A12379) for 20 minutes at room temperature. After three more PBS washes, 300 nM DAPI (Life Technologies, #D3571) in PBS was added for 10 minutes to stain nuclei. After a final PBS wash, the coverslips were mounted onto glass slides using DAKO mounting medium (DAKO Cytomation, #S3023).

### Antibodies

The following antibodies were used for immunofluorescence (IF) (Table 1):

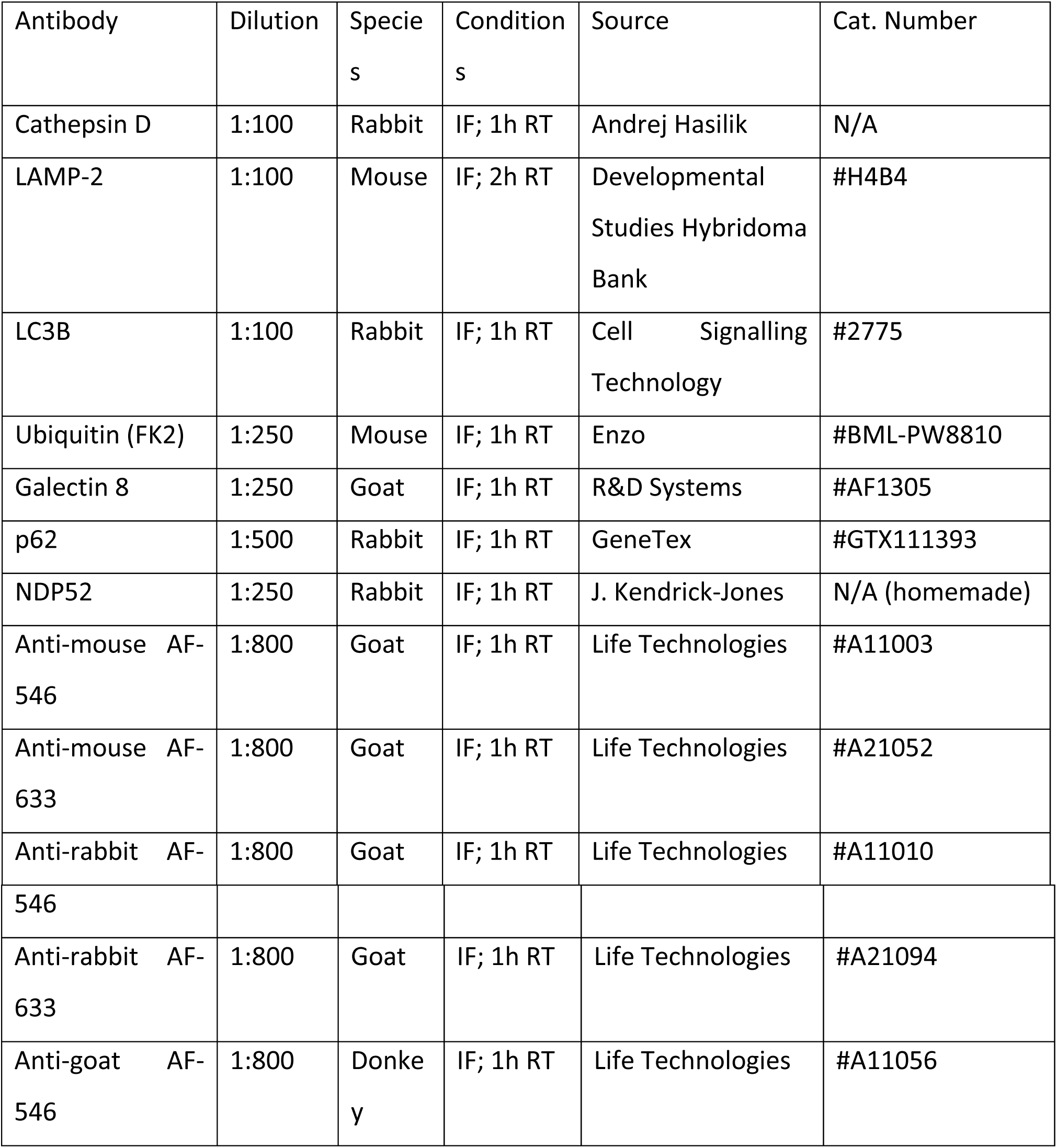

| Antibody | Dilution | Species | Conditions | Source | Cat. Number |
| --- | --- | --- | --- | --- | --- |
| Cathepsin D | 1:100 | Rabbit | IF; 1h RT | Andrej Hasilik | N/A |
| LAMP-2 | 1:100 | Mouse | IF; 2h RT | Developmental Studies Hybridoma Bank | #H4B4 |
| LC3B | 1:100 | Rabbit | IF; 1h RT | Cell Signalling Technology | #2775 |
| Ubiquitin (FK2) | 1:250 | Mouse | IF; 1h RT | Enzo | #BML-PW8810 |
| Galectin 8 | 1:250 | Goat | IF; 1h RT | R&D Systems | #AF1305 |
| p62 | 1:500 | Rabbit | IF; 1h RT | GeneTex | #GTX111393 |
| NDP52 | 1:250 | Rabbit | IF; 1h RT | J. Kendrick-Jones | N/A (homemade) |
| Anti-mouse AF-546 | 1:800 | Goat | IF; 1h RT | Life Technologies | #A11003 |
| Anti-mouse AF-633 | 1:800 | Goat | IF; 1h RT | Life Technologies | #A21052 |
| Anti-rabbit AF- | 1:800 | Goat | IF; 1h RT | Life Technologies | #A11010 |
| 546 |  |  |  |  |  |
| Anti-rabbit AF-633 | 1:800 | Goat | IF; 1h RT | Life Technologies | #A21094 |
| Anti-goat AF-546 | 1:800 | Donkey | IF; 1h RT | Life Technologies | #A11056 |

### Confocal microscope image acquisition and analysis

Imaging of fixed samples was performed using a Leica SP5 AOBS Laser Scanning Confocal Microscope (Leica Microsystems) exactly as detailed in (Lerner et al., 2016). Images were obtained in .lif format and imported into FIJI (NIH). Three parameters were measured using FIJI: a) *M. tuberculosis* growth using the total GFP signal per hLEC; b) The association of a marker (e.g. Galectin 8) to *M. tuberculosis*; c) Intracellular cord size using Feret diameter. a) and b) are extensively described in (Lerner et al., 2016), whereas Feret diameter is explained in **Supplementary Fig. 1**. All data were plotted and analysed using Microsoft Excel 2010 (Microsoft), GraphPad Prism 6 (GraphPad Software Inc.) or ggplot2 (Hadley Wickham) in R (The R Project for Statistical Computing).

### Automated confocal microscope image acquisition and analysis

After infection in a 96 wells plate, cells were fixed and stained with DAPI, and fluorescently-labelled phalloidin (conjugated with Alexa Fluor 633 or Alexa Fluor 488). Images were acquired using an automated fluorescent confocal microscope (OPERA Phenix, PerkinElmer) equipped with a 63X (NA 1.2) water lens and 405, 488, 561 and 640 nm excitation lasers. The emitted fluorescence was captured using 2 cameras associated with a set of filters covering a detection wavelength ranging from 450 to 690 nm. For each well, 30 to 35 adjoining fields containing 4 Z-stacks distant from 1µm were acquired. 10% overlap was applied between fields in order to generate a global image clustering all the fields in a single image. The maximum projection of the images was analysed using a dedicated in-built script developed using the image-analysis software Harmony 4.6 (PerkinElmer).

Cell segmentation: A local intensity detection algorithm applied on the DAPI channel was used to detect both Nuclei and cytoplasm (nuclei: maximal local intensity; cytoplasm: minimal local intensity).

Intracellular bacteria detection: A spot detection algorithm based on the GFP, RFP or Far Red channel (according to the fluorophore expressed by the bacterial strains) was applied for the detection of intracellular fluorescent-M. tuberculosis H37Rv (WT), H37Rv-ΔPDIM or H37Rv-ΔRD1. A manual threshold method, using non-infected wells, was applied to determine the background threshold. These spots were defined as region of interest (ROI) for the measurement of bacterial intensity and area in pixels. The relative bacterial load was expressed in bacterial area (pixel) per cell. The intracellular bacterial growth was quantified by the ratio of intracellular bacterial area per cell between T0 (5h.pi=uptake) and 3 days post-infection. For the quantification of the Feret diameter, the global image of the bacteria channel was exported from Harmony in .png before being converted in 8bit image and analysed in Fiji as previously described [(Lerner et al., 2016) and **Supplementary Fig. 1**].

### Live cell imaging

hLEC were seeded and infected as previously described. 24 hours prior to infection, the cells were transduced with LentiBrite RFP-p62 Lentiviral Biosensor (Merck Millipore, #17-10404) using an MOI of 40 according to the manufacturer’s instructions. After infection of hLEC, the live cell dishes were placed in a holder custom-made for confocal microscopy in a Biosafety Level 3 (BSL-3) laboratory and imaged using the following conditions: 15 min frame intervals, Z-stacks of 5 slices with 1.38 μm thickness, line averaging 4 and zoom of 1.

### Electron microscopy (EM) of single-infected cells

Electron microscopy was performed exactly as previously described (Lerner et al., 2016). Briefly, hLEC were infected for 5 + 72 hours with *M. tuberculosis* WT-EGFP prior to fixation in 4% PFA/2.5% GA in 0.1 M phosphate buffer for 24 hours at 4°C. The field of view of interest was imaged first by confocal microscopy, and then processed for imaging by serial block face scanning electron microscopy (SBF SEM) using a 3View2XP (Gatan, Pleasanton, CA) attached to a Sigma VP SEM (Zeiss, Germany). The same field of view was captured thus facilitating the creation of a composite correlative light/electron microscopy (CLEM) image. SBF SEM images were collected at 1.8 kV using the high current setting with a 20 µm aperture at 5-10 Pa chamber pressure and a 2 µs dwell time. Maximum intensity projections of confocal slices were aligned manually to highlight bacteria positions.

### Measurement of intracellular *M. tuberculosis* volumes

Selected bacteria were segmented manually from slices of SBF SEM datasets and 3D reconstructions were made using the 3dmod program of IMOD (Kremer et al., 1996). Each dataset was first de-noised with a 0.5 pixel Gaussian blur filter applied in Fiji (ImageJ; National Institutes of Health). 2 datasets from each of 2 independent samples were then segmented for each of the cord and membrane-bound bacteria conditions. The dataset xy pixels were 9.9 nm and 8.7 nm for cord bacteria, and 5.4 nm and 6.3 nm for membrane bound bacteria; all datasets consisted of serial images of 50 nm thickness. The dataset dimensions were 81.1 × 81.1 × 5.55, 71.3 × 71.36 × 1, 22.1 × 22.1 × 1.55, 51.6 × 51.6 × 2.75 µm in xyz, with 111, 20, 31, and 55 serial images, respectively. To calculate bacterial volumes, IMOD calculated Volume Inside Mesh using 3D mesh structures derived from closed contours drawn around bacteria each 50 nm, using imodmesh. For CLEM of representative 3D reconstructions of bacteria, an SBF SEM slice was assigned to a confocal slice manually in z. The confocal slice was then processed in Fiji; first, to improve interpolation during TurboReg alignment, the confocal image was upscaled from 1024 to 2048 pixels with a bilinear interpolation, and a Smooth filter applied twice; then TurboReg was then used to align the processed confocal slice with the SBF SEM image using a Scaled Rotation transformation and bacteria as landmarks (identified by fluorescence and morphology). The remaining SBF SEM images in the stack were further denoised with a 1 pixel Gaussian blur filter and brightness/contrast adjusted to match the CLEM image in Photoshop. The CLEM image was then inserted into the stack, and a Snapshot taken of the bacterial segmentation with the stack in the Model view of 3dmod.

### CLEM of co-infected cells

hLEC were co-infected with *M. tuberculosis* WT-RFP and M. tuberculosis ΔRD1-GFP prior to fixation and confocal microscopy as above. The field of interest was then processed for imaging by transmission electron microscopy (TEM). The cells were post-fixed in 1% reduced osmium tetroxide, stained with tannic acid, and quenched in 1% sodium sulphate. Next, the cells were dehydrated progressively up to 100% ethanol and incubated in a 1:1 propylene oxide/epon resin mixture. After infiltrations in pure resin, the samples were embedded at 60°C for 24 h. SBF SEM and TEM was performed as described previously (Russell et al., 2017). Briefly, the field of interest was approached by SBF SEM (there being sufficient signal for approach imaging even though the cells were not processed for this method), then the cut face was aligned to a diamond knife in a UC7 ultramicrotome (Leica Microsystems) and 70-80 nm sections from the field of interest were collected. The sections were stained with lead citrate and imaged in a TEM (Tecnai G2 Spirit BioTwin; Thermo Fisher Scientific) using a charge-coupled device camera (Orius; Gatan Inc.). For CLEM overlay, TEM images were assigned to confocal slices manually in z. The confocal slice was then processed and aligned with TurboReg in Fiji as above.

### Histology, immunohistochemistry and analysis

The study was performed using excised cervical lymph node tissue stored within the Department of Anatomical Pathology at Groote Schuur Hospital (Cape Town, South Africa). All of these biopsies were taken for clinical indications. Residual paraffin-embedded blocks of these specimens were stored for further processing. This study complied with the Declaration of Helsinki (2008), and ethics approval was obtained from the University of Cape Town Human Research Ethics Committee (REC187/2013). Informed consent was waived, as this was a retrospective study of formalin-fixed paraffin-embedded tissue samples collected during the course of routine clinical practice. Patient identifiers were unavailable to investigators.

Formalin-fixed paraffin-embedded tissue sections from patients diagnosed as tuberculosis culture positive and/or acid-fast bacilli positive (AFB+) were selected for the study and processed as described before (Lerner et al., 2016). Briefly, tissue sections were deparaffinized in xylene (2 × 10 min, 100%, 95% and 80% ethanol (2 min each). Tissue sections were then placed into an antigen retrieval buffer (Access super antigen solution, Menarini diagnostics, UK) in a decloaking chamber (Biocare Medical, CA, USA); incubated at 110 degrees for 10 min and allowed to cool for 60 min. Sections were permeabilized in PBS-0.2% Triton X-100 and incubated in blocking buffer (1% BSA, 5% Fetal Calf Serum in PBS) overnight at room temperature. Primary and secondary antibodies were tested for cross reaction in samples of uninfected individuals. Primary (human antigens) and secondary antibodies for cross-reaction with *M. tuberculosis* in samples that were acid fast positive (AFB+).

### RNA extraction and sequencing library preparation

*M. tuberculosis*-infected or uninfected hLECs were lysed in 0.5mL of TRIzol and RNA was extracted using Direct-zol RNA MiniPrep Kit (Zymo Research) and treated with TURBO DNase I (Life Technologies) until DNA-free. Quantity and quality of the extracted RNA were determined by Qubit flourometer, NanoDrop spectrophotometer and Bioanalzyer. RNA-Seq libraries were prepared using 1mg of RNA of each sample with TruSeq Stranded Total RNA Library Prep kit (Illumina) and ribosomal RNA was removed with Ribo-Zero as part of the library construction process. Quality and quantity of the cDNA libraries were determined by Qubit flourometer and Bioanalzyer before being processed for sequencing with Illumina Hi-Seq 2500 for single-end reads with 100 cycles.

### RNA-Seq data analysis

The RNA-Seq data in this paper have been deposited in Gene Expression Omnibus repository with accession number **GSE110564**. The quality of the Illumina-produced fastq files was assessed using FastQC (v0.11.5) and adapter trimmed using Trimmomatic (v0.36). The resulting reads were then aligned to the human genome (Ensembl GRCh38 release 88 build) using STAR aligner (v2.5.2a). Gene counting was done using RSEM (v1.2.29) and expected read counts were normalized using DESeq2 (v1.18.1), which also determined the log2 fold change and statistical significance between the infected and uninfected samples. Canonical pathway and functional process analyses were performed using IPA Ingenuity (QIAGEN) and MetaCore (Thomson Reuters).

### Real-time polymerase chain reaction (RT qPCR)

Isolated RNA was processed with QuantiTect™ Reverse Transcription Kit (Qiagen). Quantitative real-time RT–PCR (qRT–PCR) was performed using 11.25 ng cDNA per well with 0.5 μl TaqMan™ Gene Expression Assay probe and 5 μl TaqMan™Universal PCR Master Mix in a 10-μl reaction volume on The Applied Biosystems™ QuantStudio™ 7 Flex Real-Time PCR System. Each reaction was performed in triplicate. Data analysis was performed using ExpressionSuite for QuantStudio™ (Applied Biosystems). Fold change was determined in relative quantification units using GAPDH for normalization.

### Data and statistical analysis

Results are expressed as mean ± SEM. All statistical analyses were performed in Prism 6 (GraphPad Software Inc.). Means between 2 groups were compared using two-tailed Student’s *t* tests and means among 3 or more groups were compared using one-way ANOVA with Tukey’s multiple comparisons tests. A *p* value of under 0.05 was considered significant (*p<0.05; ** p<0.01, *** p<0.001). Plots were produced in Prism 6 or ggplot2 in R (The R Project for Statistical Computing).

## Author contributions

MGG and TRL conceived the project. MGG, TRL, CJQ and RPL designed the experiments. TRL, CJQ, RPL, MGR, AF and CQJ performed experiments. TRL, CJQ, RPL, MGR, AF, LC, DJG and RJW analysed data and provided intellectual input. MGG wrote the manuscript with input from TRL. All authors read the manuscript and provided critical feedback.

## Acknowledgements

We thank J. Kendrick-Jones (MRC-LMB, Cambridge) for NDP52 antibody and Michael Niederweis (University of Alabama) for fluorescent plasmids. Bill Jacobs (Albert Einstein College of Medicine), Suzie Hingley-Wilson (University of Surrey), Catherine Astaire-Dequeuer (IPBS, Toulouse), Douglas Young (Crick) and Sebastien Gagneux (THP, Basel) for *M. tuberculosis* strains, Steve Coade for assistance with fluorescent tagging of clinical isolates and Susanne Herbst for critical reading of the manuscript. This work was supported by the Francis Crick Institute (to MGG and RJW), which receives its core funding from Cancer Research UK (FC001092, FC00110218), the UK Medical Research Council (FC001092, FC00110218), and the Wellcome Trust (FC001092, FC00110218) and Wellcome Trust (to RJW, 104803, 203135).

## Supplementary figure legends

**Supplementary Fig. 1:**
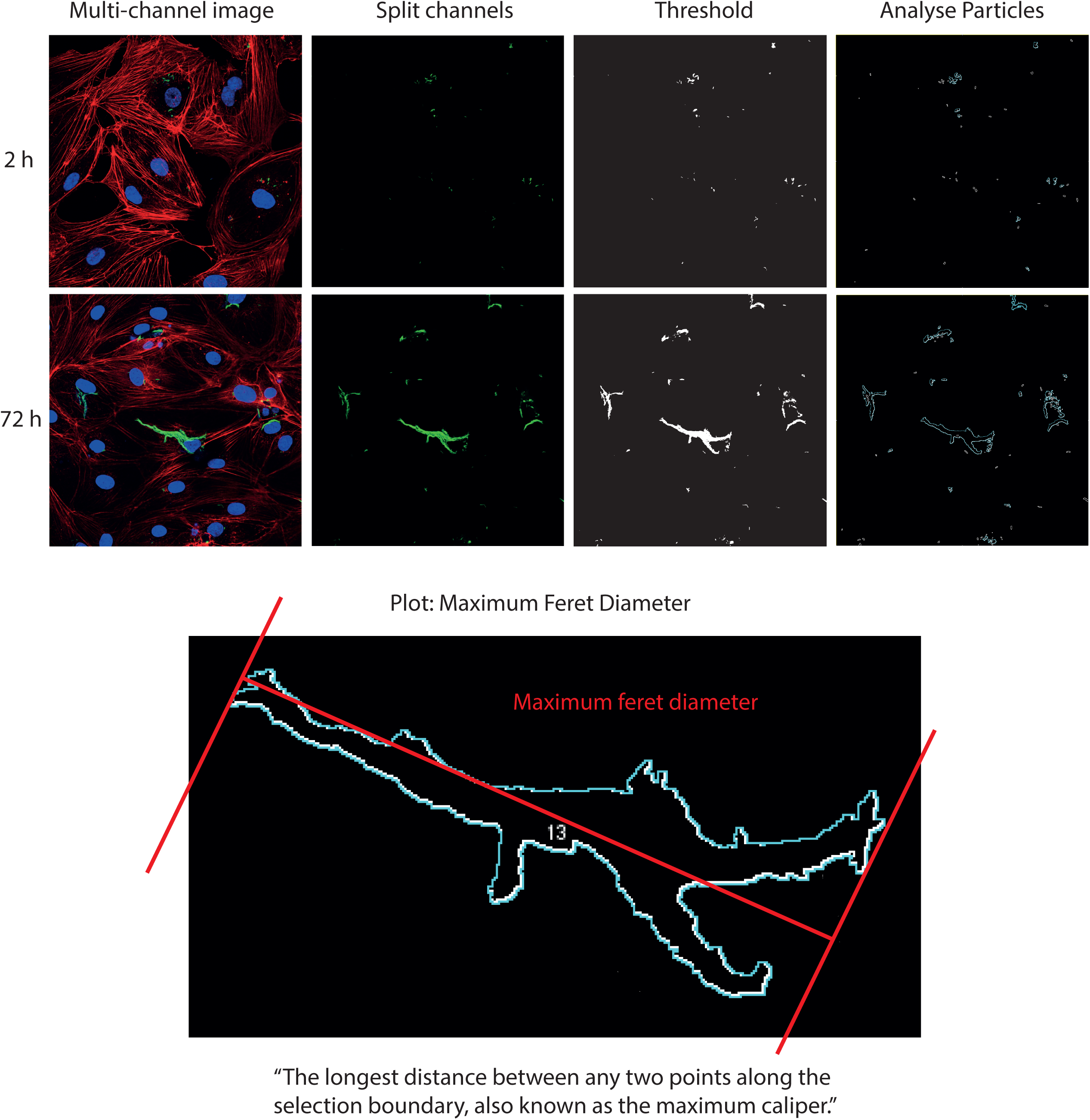
Demonstration of feret diameter as a measure of cord length. ImageJ was used to first select only the GFP channel corresponding to the bacteria. The GFP was subjected to a pixel threshold, dilated and eroded (add and then remove 1 pixel to the outline, to link up any incorrectly thresholded pixels) and then outlined to form ‘particles’ which were analysed by Feret diameter. The feret diameter is calculated as shown; it is the distance between the two furthest apart pixels in the particle.

**Supplementary Fig. 2:**
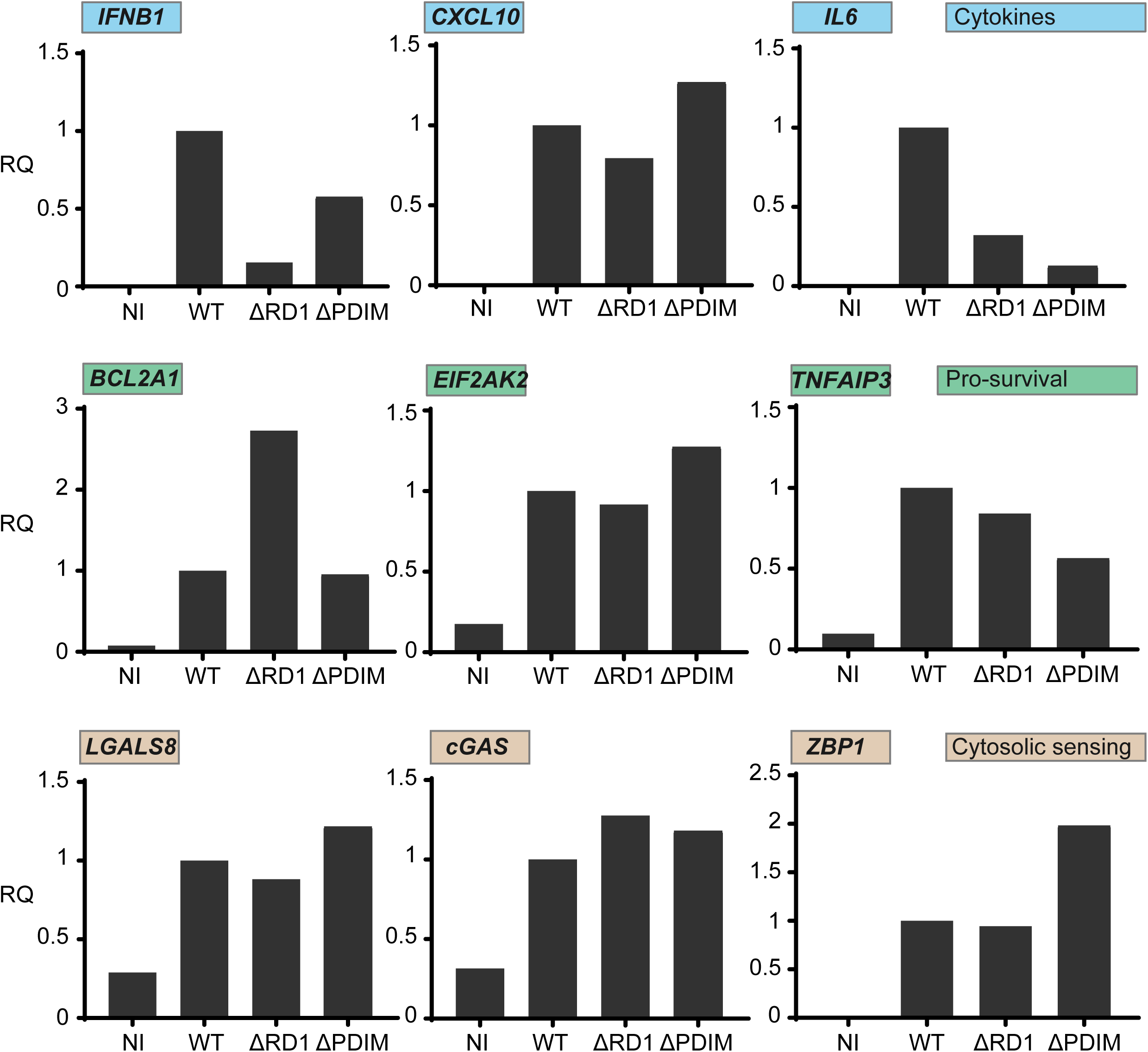
RD1 and PDIM-dependent regulation of selected gene expression in hLEC after infection. Gene expression measured by RT-qPCR of infection pathways, displayed in Fig 2, in response to hLEC infection with *M. tuberculosis*-H37Rv WT, *M. tuberculosis*-H37Rv ΔRD1 or ΔPDIM for 48 h post infection. Uninfected hLEC were used as control of the basal expression of each gene. Data are representative of 2 independent experiments, each carried out in duplicate.

**Supplementary Fig. 3:**
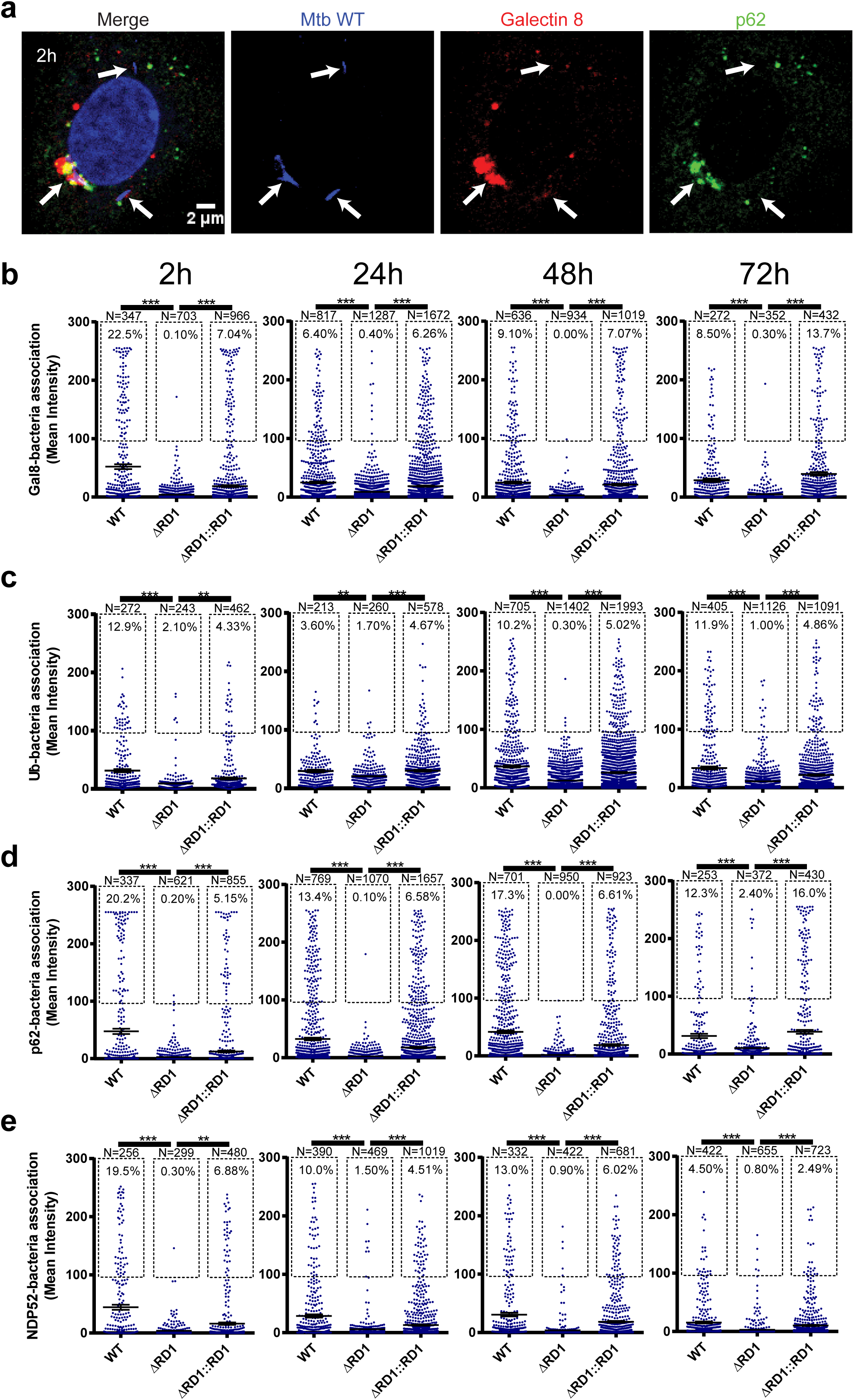
*M. tuberculosis* targeting via selective autophagy in hLECs is RD1 dependent. **(a)** hLEC were infected with *M. tuberculosis* WT-GFP (blue) and stained with galectin 8 (red) and p62 (green). Nuclei were stained with DAPI (blue). White arrows show the localisation of *M. tuberculosis* in each channel. Scale bar is 2 µm. **(b-e)** hLEC were infected with *M. tuberculosis* WT, *M. tuberculosis* ΔRD1 or *M. tuberculosis* ΔRD1::RD1 (all GFP tagged) for 2, 24, 48 or 72 h before fixation. The samples were labelled using antibodies against **(b)** Galectin 8 (Gal8), **(c)** Ubiquitin (Ub), **(d)** p62 or nuclear dot protein 52 (NDP52). Nuclei were visualised with DAPI. Images such as in **(a)** were quantified using ImageJ to measure the association of each marker to each strain of *M. tuberculosis* at each time-point. The percentage of *M. tuberculosis* that were positive for the marker (defined as the marker (the red channel) having a mean pixel intensity of at least 100 in the area overlapping with the individual *M. tuberculosis* particle (the green channel)) in each condition is shown. At least six fields of view were imaged per sample per replicate and three independent replicates were performed. The total number of analysed *M. tuberculosis* particles is also displayed for each condition (N). The overall mean is shown with error bars representing the standard error of the mean (SEM). Statistical significance was determined using One way ANOVA with Tukey’s post-test; *** = p < 0.001.

## Movie S1

Human lymphatic endothelial cells (hLEC) expressing p62-RFP (red) infected with *M. tuberculosis* expressing EGFP (green) were imaged for 100 h. Left hand side shows an overlay of phase contrast and fluorescence images, whereas right hand side shows fluorescence images only. The purple arrows track an example of an individual *M. tuberculosis* bacterium that does not become p62 positive at any point and eventually grows into a long intracellular cord. Subsequently, the infected cell dies and detaches. The blue arrows track an example of *M. tuberculosis* that becomes p62 positive soon after uptake, remains p62 positive, and shows severely restricted growth.

## Movie S2 and S3

Similar to Movie S1 except the blue arrows track an example of *M. tuberculosis* that fluctuates between having p62 association and not. Eventually, once p62 association ceases, an intracellular cord forms.

